# Erroneous inference based on a lack of preference within one group: autism, mice, and the Social Approach Task

**DOI:** 10.1101/530279

**Authors:** Kayla R. Nygaard, Susan E. Maloney, Joseph D. Dougherty

## Abstract

The Social Approach Task is commonly used to identify sociability deficits when modeling liability factors for autism spectrum disorder (ASD) in mice. It was developed to expand upon assays available to examine distinct aspects of social behavior in rodents and has become a standard component of mouse ASD-relevant phenotyping pipelines. However, there is variability in the statistical analysis and interpretation of results from this task. A common analytical approach is to conduct within-group comparisons only, and then interpret a difference in *significance* levels as if it were a group difference, without any direct comparison. As an efficient shorthand, we named this approach **EWOCs:** <u>E</u>rroneous <u>W</u>ithin-group <u>O</u>nly <u>C</u>omparison<u>s</u>. Here we examined the prevalence of EWOCs and used simulations to test whether it could produce misleading inferences. Our review of Social Approach studies of high-confidence ASD genes revealed 45% of papers sampled used only this analytical approach. Through simulations, we then demonstrate how a lack of significant difference within one group often doesn’t correspond to a significant difference between groups, and show this erroneous interpretation increases the rate of false positives up to 25%. Finally, we define a simple solution: use an index, like a social preference score, with direct statistical comparisons between groups to identify significant differences. We also provide power calculations to guide sample size in future studies. Overall, elimination of EWOCs and adoption of direct comparisons should result in more accurate, reliable, and reproducible data interpretations from the Social Approach Task across ASD liability models.

**Lay Summary:** The Social Approach Task is widely used to assess social behavior in mice and is frequently used in studies modeling autism. However, reviewing published studies showed nearly half do not use correct comparisons to interpret the data. Using simulated and original data, we argue the correct statistical approach is a direct comparison of scores between groups. This simple solution should reduce false positives and improve consistency of results across studies.

## Introduction

The Social Approach Task is one of the most widely utilized behavioral assays for investigation of mouse models of liability factors associated with autism spectrum disorders (ASD) (Bader et al., 2011; Chadman et al., 2008; Copping et al., 2017; Dougherty et al., 2013; Feyder et al., 2010; Grabrucker, Boeckers, & Grabrucker, 2016; Lugo, Swann, & Anderson, 2014; Maloney et al., 2018; Page, Kuti, Prestia, & Sur, 2009; Penagarikano et al., 2015; Samaco et al., 2012; Schwartzer et al., 2013; Stoppel et al., 2018; Won et al., 2012) The use of the Social Approach Task has both helped identify ASD liability models with good face validity, and advanced our understanding of the circuitry underlying social approach deficits. Unlike reciprocal social interaction assays requiring manual scoring, the Social Approach Task is automated, making it ideal for mechanistic studies that require several experiments with different interventions or genetic models. For example, a recent study showed the role of dorsal raphe serotonergic connections to the nucleus accumbens in social approach behavior, and how stimulation of this pathway can correct social deficits in the 16pll.2 deletion model associated with ASD (Walsh et al., 2018). Another group showed NMDAR activation rescued social approach behavior in *Shank*^-/-^ and *Tbr1*^+/-^ mutants (Lee et al., 2015; Won et al., 2012). Together these studies show the value in using this task to identify pathways that contribute to social approach behavior and targets that can be further interrogated as potential pharmacotherapy candidates.

The motivation behind development of the Social Approach Task was to improve face validity of murine social behavioral assays with regards to specific social impairments that characterize ASD (Moy et al., 2004; Nadler et al., 2004). Abnormal social approach is one such attribute of the ASD social phenotype. This task was unique in the field because it required the sociability be initiated by the test mouse. Thus, it was, and is, meant to help identify a lack of social interest in mice that may be reminiscent of the social approach deficits in humans with ASD. This task comprises two test trials: the *sociability trial* and *preference for social novelty* trial (Moy et al., 2004; Nadler et al., 2004). During the *sociability* trial, the test mouse can freely explore the 3-chambered apparatus to investigate either a novel conspecific stimulus in a restraining container (inverted wire cup) or an empty but otherwise identical wire cup (Figure 1A). If only one group of mice is being evaluated, a comparison between time spent with the empty cup and time spent with the social cup (novel conspecific) can be used to demonstrate this group has a significant preference for social stimuli, as is observed with wild-type (WT) mice of most strains (Moy et al., 2004). However, since the task was first developed, the within-group only analysis has also been perpetuated across studies of between-group factors, such as ASD candidate genes.

**Figure 1:**
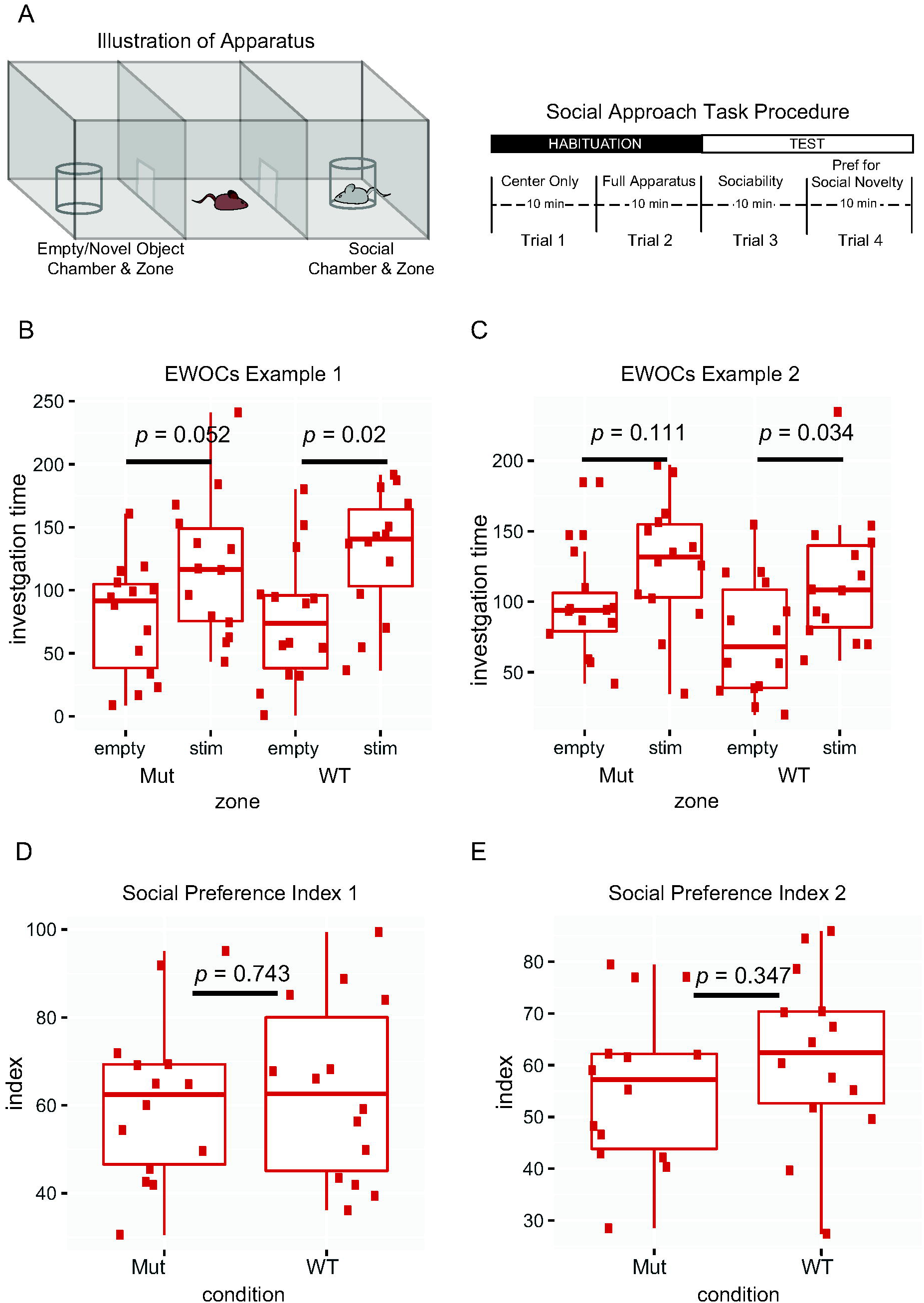
Illustration of the Social Approach Task and two different analytical approaches. **A)** Schematic of Social Approach Task and apparatus. **B,C)** Example plots from simulated data using EWOCs. Two arbitrary groups (‘Mut’ and ‘WT’) were tested for a within-group difference between the time spent with the social stimulus (stim) compared to the empty cup (empty). Only the WT group showed significant preference (*p*<0.05), while the Mut mice did not (*p*=0.052, orp=.lll). **D,E)** Example of the same data plotted as a social preference index 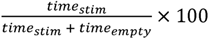. statistical comparison of Mut to WT indices shows no significant difference (*p*=0.743, 347).

While a within-group analysis is necessary to determine that a control group has performed as expected and to support the assay’s validity, after careful consideration of this task, it occurred to us the fairly standard interpretations derived from <u>only</u> using within-group comparisons of time spent with social versus empty stimuli across multiple groups may be statistically and logically inappropriate. Here, we present reasons for why <u>E</u>rroneous <u>W</u>ithin-group <u>O</u>nly <u>C</u>omparisons (EWOCs) should not be applied to identify a difference between groups. We argue both from the perspective of statistical principles and by using data simulation to show how EWOCs can be misleading. We also review recent mouse literature to characterize the use of EWOCs. We further show how direct comparison of an index, like a social preference score, between groups may reduce false positives and improve consistency of results across studies, and provide power estimates, parameterized in data from >400 mice, to guide future studies. We believe elimination of EWOCs from practice will result in more robust and reproducible social approach findings when modeling ASD liability factors in mice.

## Methods

### Simulation studies

We conducted multiple analyses of simulated data to explore the frequency of erroneous inferences when using only EWOCs to determine a difference between groups. First, we collected all data previously generated for social approach in the lab, which includes 231 mice from (Dougherty et al., 2013; Maloney et al., 2018) and an additional 190 mice subsequently tested. Using this data, we calculated the mean interaction time in seconds (s) with the stimulus mouse (time_[stim]_; 124.06 ±52.90 [SD]), and the mean time with the empty cup (time_[empty]_; 87.51 ±40.59 [SD]). We then wrote a simple function in R to generate 1000 random experiments with a sample size of 10 per group using the function *rnorm* to sample two arbitrary groups (‘Mut’ and ‘WT’) from the same normal distribution with parameters derived from data above (timeS_[stim]_ =124s, time_[empty]_ =88s, *SD* =47s). Using this function, we calculated the frequency of incorrect interpretations when utilizing EWOCs (conducting separate t-tests comparing time_[stim]_ to time_[empty]_ for Mut and WT groups and comparing the results) and repeated the thousand-experiment simulation ten times. Second, we repeated this method and systematically varied the group sample size *(n)* from 2 to 30 to illustrate the vulnerability of EWOCs to false positives across *n,* and what happens when *n* is mismatched between groups. Third, we modeled the consequences of varying the magnitude of social preference by changing the mean of the sampled normal distributions across a range of values. We set indices for a range of social preference values from 50 (no preference) to 75 (a 3-fold preference for the stimulus mouse) by setting values of time_[stim]_ from 106 to 159 s (and correspondingly adjusted the mean for time_[empty]_). Fourth, we modeled the effect of differential group variability by increasing the standard deviation of only the Mut group from 47 to 78 but keeping the mean preferences the same for both groups. We then repeated all the above analyses, but instead calculated the frequency of erroneous inferences when transforming the time into a social preference index, defined as 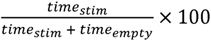, conducting a *t*-test comparing indices of the two groups. In addition, we duplicated all our analyses using simulations based on parameters extracted from a published paper (Filipello et al., 2018), using chamber time instead of investigation zone time, which yielded substantially similar conclusions.

### Systematic review of the literature

To assess the potential impact of EWOCs in ASD-related research, we performed a systematic review of the literature, utilizing the SFARI Animal Models database (Kumar et al., 2011) (accessed July 18, 2018) to identify relevant literature. Focusing on only genes with a score of 1, which indicates “High Confidence” autism genes, we reviewed all referenced papers for each gene to identify studies using the 3-chambered Social Approach Task. These studies were used for the next round of analysis where we extracted the results for the *sociability* and *preference for social novelty* trials, sample size, and whether EWOCs were used. If a study used both within-group and between-group comparisons, it was not counted as an EWOCs study. Finally, an independent researcher reread all studies to confirm only this interpretation was used.

### Power calculations

We estimated the required group sizes using the pwr.t.test function in R, using settings for two samples with a one-tailed (“greater than”) hypothesis. We ran the algorithm for three magnitudes of power (.7, .8, and .9) and systematically varied the effect size across a range of plausible values. We parameterized our calculation of effect size (Cohen’s *d)* using values based on the >420 mice from our lab. Specifically, we set the pooled standard deviation for the social preference index at 15.64 (the standard deviation of our mice), calculated effect sizes assuming a mutant group would have no social preference (a group mean of 50) and varied the corresponding wild-type preference to between 54 (below the group mean of FVB mice at 54.81) to 66, slightly above our most social group: C57BL6/J males (63.3), and the mean preference of the reviewed published studies (64.17). Resulting group sizes were then plotted as function of effect size and desired power.

## Results

### Interpreting EWOCs as a difference between groups is fundamentally flawed logic

The experimental hypothesis of the traditional Social Approach Task is the test subject will spend more time with a social stimulus relative to a nonsocial stimulus, which is interpreted as a social preference. To test statistical significance of the experimental results, data are compared with a null hypothesis, in this case, ‘the test subject will spend equal time with both stimuli’. When results are interpreted as significant, the null hypothesis is rejected but the experimental hypothesis is not accepted. To avoid false positives, the cutoff or critical probability level (alpha) is lowered, and to avoid false negatives, the sample size *(n),* and thus corresponding statistical power, is increased.

While it is relatively straightforward to test significance in the Social Approach Task within one group, a variety of approaches have been utilized when comparing two different groups. Specifically, many papers do not use identical methods to compare the social preferences of a WT control with those of a disease model. Often, social preference is tested within each group, but not between. Thus, the lack of a statistically significant preference in one group is interpreted as a statistically significant difference between groups. This is a flawed interpretation because the null hypothesis is no longer accurately tested. The new null hypothesis when comparing multiple groups is ‘the social preference of one group equals the other’. This requires a direct test between the groups, which can be done using a repeated measures ANOVA with appropriate between-subjects factors to examine stimulus interaction times. An equally valid approach is to calculate a single value summarizing social preference for each mouse for downstream statistical testing. A commonly used social preference index is 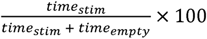, which results in a value of 50 (equal time with both) to 100 (all time with stimulus mouse). Indices for each mouse can then be compared across groups with a *t-*test, ANOVA, or appropriate non-parametric test for nonnormal data.

To illustrate how a within-group only comparative approach to analysis could lead to erroneous inference, consider the following example (Figure 1B). In this simulated data of a *sociability* trial, the mutant mice do not show a statistically significant social preference, with a p-value=0.052. As this exceeds the critical alpha cutoff of 0.05, it does not result in a rejection of the null hypothesis. The WT mice, however, reach a p-value=0.02, which passes the cutoff. The null hypothesis is rejected, and the WT mice are considered to have shown a statistically significant preference for the social stimulus. Even though the outcome of the tests within the groups are different for mutant and WT mice, does this mean there is significant difference in the social preference between these groups? In this example, where p-values are just on either side of the threshold, it becomes obvious that a separate statistical test is necessary to determine if the groups themselves are statistically different. Indeed, calculating a social preference index and comparing them directly for this same data reveals there is no difference between the groups (Figure 1D). To reiterate, a lack of difference within one group does not indicate a significant difference between groups.

Unfortunately, this simple statistical misinterpretation exists widely in the neuroscience literature and is applied to many kinds of experiments (Nieuwenhuis, Forstmann, & Wagenmakers, 2011). It also exists in key papers evaluating genetic mouse models of ASD liability. To assess the impact of EWOCs, we systematically reviewed the mouse literature referenced in the SFARI database for genes with a score of 1, classified as High Confidence. We further limited this to the 29 papers that used the Social Approach Task, including both the *sociability* and *preference for social novelty* trials. Looking at the mutant data, across these studies, EWOCs were employed in 13 of 29 (44.8%) studies showing a phenotype in the *sociability* trial, and 11 of 25 (44.0%) of studies for the *preference for social novelty* trial. Thus, use of EWOCs are widespread.

This raises important questions: To what extent might these represent false positive results? Could widespread use of EWOCs account for why there are such challenges in finding reproducible phenotypes in behavioral models (Kafkafi et al., 2018)? In order to determine how vulnerable this approach is to false positive interpretations, we conducted extensive simulation studies as detailed below.

### Simulation demonstrates EWOCs result in an elevated rate of false positives, dependent on sample number

We first modeled how likely false positive results would be when using EWOCs. To base the simulation on real parameters, we examined social approach data from all mice previously tested in our lab to identify typical mean interaction times and standard deviations. We also extracted the data examined in all 29 datasets from the reviewed papers for comparison. We found the median group size was *n ≈* 16 across the 29 papers (Figure 2A), with studies ranging from 6 to 30. We then generated random data for two groups with no true difference in their social preference (drawing from the same normal distribution) such that both ‘WT’ and ‘Mut’ groups should have a 1.5-fold preference for the social stimulus over the empty cup (social preference index=60; Figure 2B). We then systematically varied the *n* in each group from 5 to 30 and conducted 10 simulations of 1000 studies at each *n.* When we simulated *n* at the median of published studies (i.e. 16 per group), we observed a false positive rate of 25% using EWOCs (Figure 2C). Even extending *n* to 25, we still observed a false positive rate of 10%, which is approximately 2 times higher than the false positive rate of 0.05 that is the standard accepted critical alpha in the field. Note that a solution for controlling the false positive rate is quite simple: a t-test assessing the social preference index, with *p* < 0.05 as a cutoff, results in the expectedly well-controlled false positive rate of 5%, regardless of *n* (Figure 2D). Similarly well-controlled results are also achieved if one analyzes the stimulus interaction times across groups using a mixed ANOVA with between- and within-subjects simple main effects following significant interaction terms *(not shown).* Importantly, if *n* is imbalanced, then statistical power is also imbalanced. For example, sometimes mutants are harder to generate than WT (indeed, 1/3 of the reviewed studies had smaller mutant than WT groups). This might further inflate false positive rates when EWOCs are used. By varying *n* for ‘Mut’ with n=12 for ‘WT,’ we show this is the case (Figure 2E). Again, this can be corrected by directly comparing groups statistically (Figure 2F).

**Figure 2:**
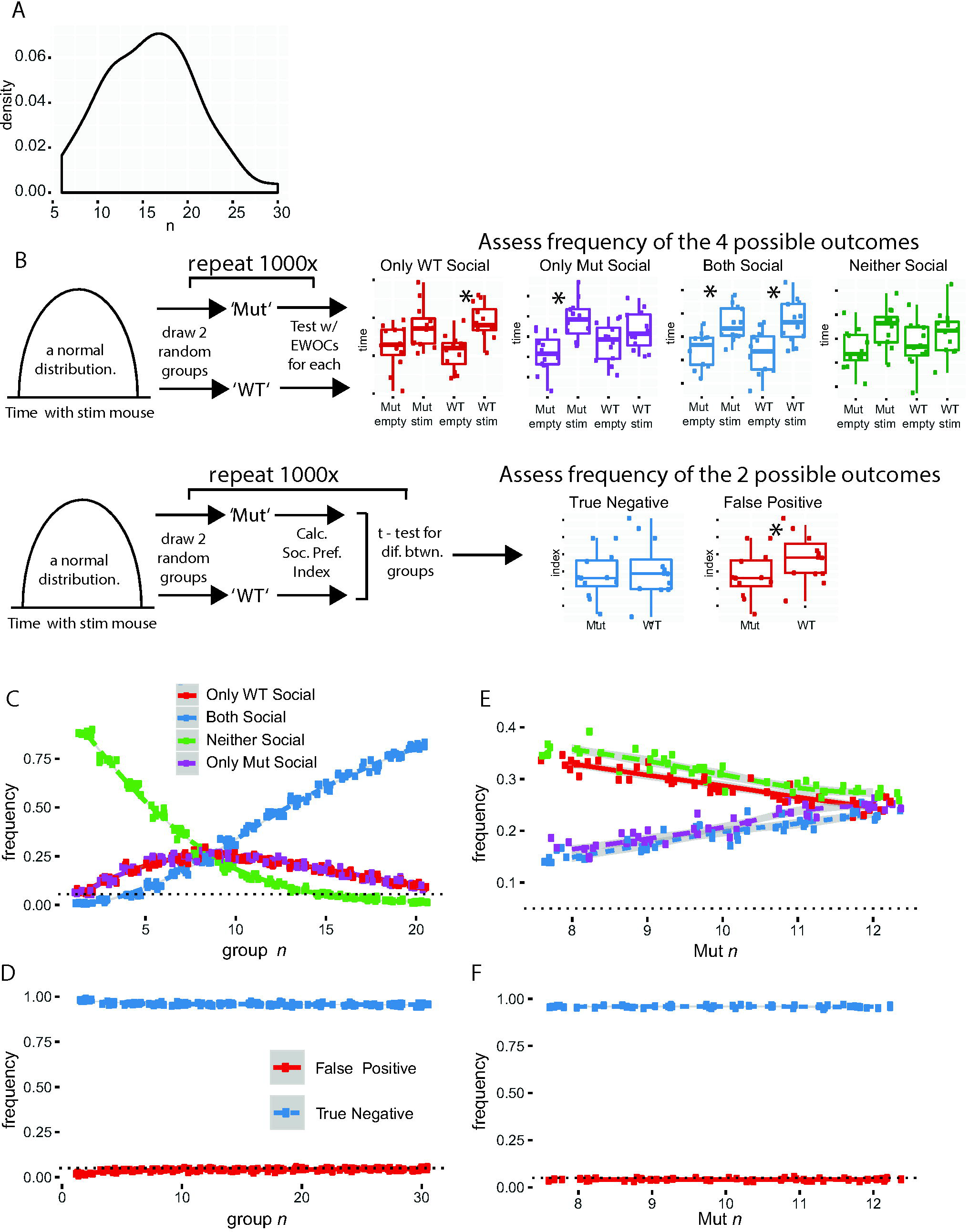
using EWOCs can result in substantially elevated false positive rates, especially at low sample sizes. **A)** Distribution of group sizes (combined for genotype) across 77 groups in 29 papers. **B)** Cartoon of simulations and possible outcomes. Two groups (‘Mut’ and ‘WT’) are drawn from the same distribution with identical social preference magnitude, and then tested with EWOCs *(upper panel)* or a social preference index *(lower panel).* **C)** Plot of simulations results as function of *n,* after 10 × 1000 simulated experiments for each *n,* drawing two groups from the same distribution and analyzing with EWOCs. The true result is both groups are social *(Blue),* so incorrect conclusions were drawn a substantial proportion of the time. **D)** Plot of *t*-test on social preference index, showing false positive rate as a function of *n.* **E)** Simulation plot as a function of imbalanced *n* with WT *n*=12, and Mut *n* varied from 8 to 12, using EWOCs. **F)** Simulation plot as a function of unbalanced *n,* using *t*-test on the social preference index.

It is worth noting that even with equal *n,* other results can also occur. For example, if WT and mutant mice are truly not different, there is an equal chance that the ‘Mut’ mice will show a significant preference for the social stimulus in the same trial that the ‘WT’ mice do not (Figure 2B,C; purple lines). There is also a chance, especially at low *n,* that neither group will show a significant within-group result (Figure 2B,C; green lines). Given the known bias in published literature for positive over negative results (Matosin, Frank, Engel, Lum, & Newell, 2014), it is likely that either of these possibilities are underreported in the literature. For example, they may simply be considered failed trials by the experimenters and repeated, since the positive control (i.e. a preference for the stimulus mouse in the WT group) did not work. One danger of this repeated EWOCs approach is that it could further increase the possibility of a false positive, as the experiment would be repeated until the outcome is either both groups are social, or only the mutants have a deficit. Overall, even with a single experiment of simulated data at *n*=16, there is only a <70% chance of correctly identifying both groups as social.

### Simulation demonstrates EWOCs false positive rates are also influenced by magnitude of social preference

Of course, statistical power is also a function of effect size - in this case, the magnitude of the social preference. In our first model, we assumed a 1.5-fold preference for the stimulus mouse over the empty cage, modeling a normal distribution with a mean interaction time of 126 seconds with the stimulus mouse and 86 seconds with the empty cup (giving a social preference index of 60). While this is a plausible social preference magnitude, and slightly higher than the mean we saw in our reanalyzed mice (124.06), it is a bit below the median social preference index of published groups (64.41 [58.96-69.70 interquartile range (IQR)]; across all 77 groups of extractable data from the 29 studies; Figure 3A). Therefore, we also fixed *n* at 10 and varied the simulated preference of all mice for the social stimulus. This showed a high rate of erroneous inference resulting from EWOCs. Interestingly, a social preference index around 64 was particularly vulnerable to EWOCs false positive interpretation (Figure 3B), with rates at nearly 25%. Note, differences in effect size are also readily controlled by appropriately comparing the two groups statistically (Figure 3C).

**Figure 3:**
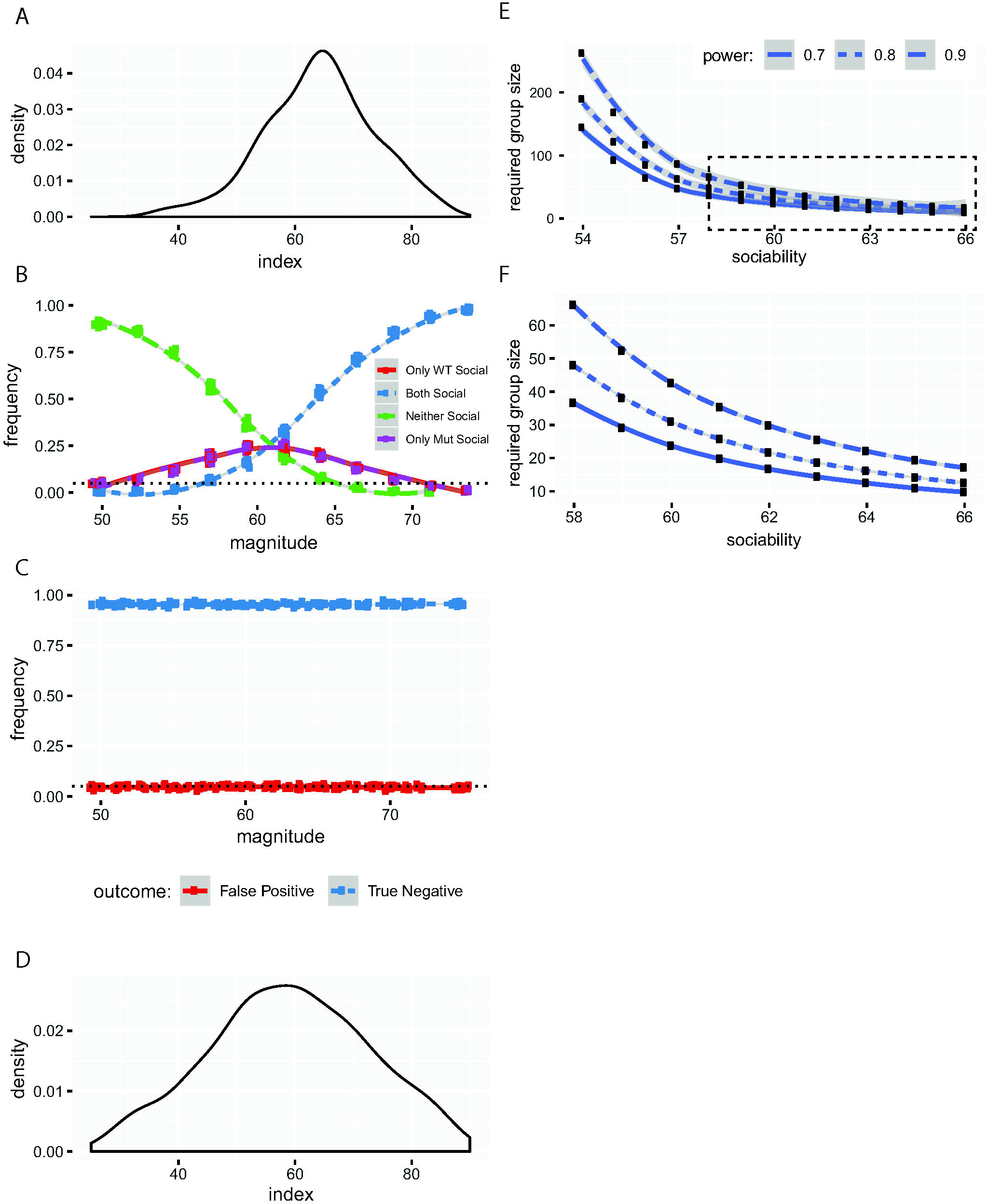
Elevation of false positive rates depends on the magnitude of the social preference when EWOCs are used. A) Distributions of average magnitudes of social preference indices across groups from the 29 reviewed studies. **B)** Plot of outcomes as a function of social preference magnitude when using EWOCs. **C)** Plot false positive rate as a function of social preference magnitude when using *t-*test on social preference index. **D)** Distributions of magnitudes of social preference indices from all mice run in our lab (*n*=421). E) Power calculations showing required *n* per group as a function of the WT social preference index, to have 70%, 80%, or 90% power to detect a difference at 0.05. F) Same, replotting boxed region from E.

Also worth discussion is the possibility the published median social preference magnitude is slightly inflated compared to the actual social preference, again, because of the bias towards publication of positive results. Indeed, if we plot the social preference index of the last 421 mice analyzed in our lab (Figure 3D), published or not, we see a median preference of 58.95 (48.95-68.48 IQR) for the *sociability* trial, and 63.49 (51.69-71.64 IQR) for the mice that were also tested in the preference for *social novelty* trial *(n=*325, *not shown).* We also noticed a commonly used inbred strain (FVB/AntJ, e.g the standard strain of FMRP mutants) showed a marginally lower social preference index than the more ubiquitous inbred C57BL/6J strain (54.8 vs. 60.1, Welch’s t-test *t*=2.3128, *p*=0.023, *df=*107.03), and, generally, males showed a higher social preference index than females across strains (60.98 vs 55.04, *t*=3.9615 *p*=8.7E-5, *df=*418.72). Thus, the expected magnitude of social preference in this task may vary by sex and strain, and may be low enough to warrant increased *n* when utilizing both sexes for experiments (as currently required by NIH funding).

Therefore, as a resource, we have estimated the number of animals required to have well-powered studies detecting an absence of social preference (i.e. social preference index of 50) in a mutant group compared to a variety of potential wild-type group preference index levels. Our estimates show that to have 80% power to detect a significant effect requires approximately 30 animals per group using both sexes of C57BL6/J mice, and possibly substantially more on other strains (Figure 3E-G), though these strains may be better when assaying manipulations that increase sociability. Further, *social novelty* trials, where the effect size is typically somewhat larger, would require less animals.

### Simulation demonstrates that behavioral disruptions that increase variance in mutants will also lead to higher false positive rates with EWOCs

Finally, there are even more subtle features of mouse behavior that might lead to inflated false positive rates with EWOCs. This is because commonly used test statistics are defined as the difference in the means divided by a measure of variance. Thus, if one group is significantly *more* variable than another, it is *less* likely to have a large test statistic and thus *less* likely to achieve a significant p-value. For example, if mutant mice tend to have a compulsive grooming phenotype making their movement in the task more stochastic (i.e. they might spontaneously enter a long bout of compulsive grooming) then their variance might simply be higher in this task compared to controls. It is hard to determine how frequently such a thing might be occurring in the literature, but it is straightforward to model – holding a constant *n* (10) and social preference index (60), we altered the variance of the distribution from which we drew the ‘Mut’, but not the ‘WT’, group. This profoundly decreased the ability to detect a significant social preference in the ‘Mut’ group (Figure 4A), and, interestingly, this phenomenon could not be readily rescued by increasing *n* (Figure 4B,C). Thus, mutations that increase variability in mouse behavior, when using EWOCs, can mask true social preference. Again, when you directly compare groups statistically, the false positive rate stays at a well-controlled 5% (Figure 4D).

**Figure 4:**
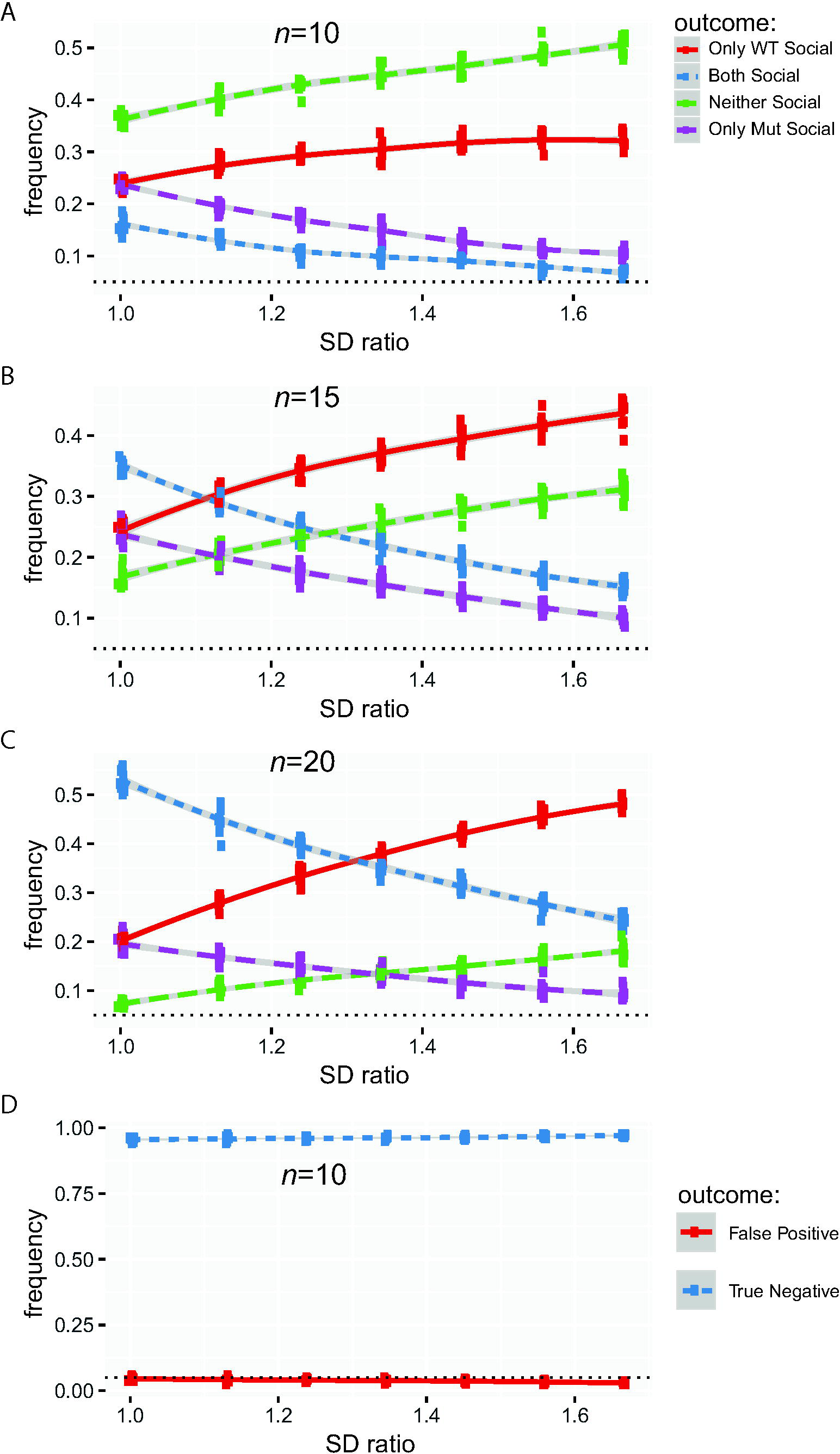
Increased variance in mutants can also lead to inflated false positive rates when EWOCs are used. **A)** Plot of false positive results when using EWOCs as a function of increased variance in only Mut at *n*=10, **B)** at *n*=15, **C)** at *n=*20. **D)** Plot of *t*-test false positive rate as a function of increased variance at *n*=10. *SD* Ratio: the ratio of the Mut to the WT standard deviation (SD; varied from 1 to 1.5).

To demonstrate that the flawed logic of EWOCs extend to chamber time data, as well, we duplicated all our above analyses using simulations based on means and standard deviations extracted from a published paper (Filipello et al., 2018) using chamber time instead of investigation zone time. The results were substantially similar *(data not shown).*

## Discussion

The Social Approach Task has been heavily relied on to assess social behavior phenotypes in genetic liability factors for ASD. Thus, it is essential to use appropriate statistical approaches to ensure proper interpretation of the results. Only this will allow for correct conclusions to be drawn about the influence of ASD candidate genes and other liability factors on social approach circuits.

In almost half of published papers based on our sampling, the interpretation of results of this task were based on within-group only comparisons without a direct comparison between the experimental and control groups. Thus, <u>E</u>rroneous <u>W</u>ithin-group <u>O</u>nly <u>C</u>omparison<u>s</u> (EWOCs) is frequently interpreted as a difference between groups. The problem with using this approach, essentially concluding that ‘if the result is not significant, sociability is absent’, is that statistical tests are designed only to identify significant differences. They are *not* designed to identify a significant *lack* of differences. In other words, the correct interpretation when p>.05 is not “We are 95% confident there is no difference in preference between the mouse and the cup.“ It is “We are *not* 95% confident that there *is* a difference between the mouse and the cup.” Statistical tests would have to be completely redesigned to be able to state with 95% confidence that there is no preference, and it is far simpler to directly compare the relevant groups with standard tests. We refer the reader back to the example in Figure 1B illustrating how EWOCs do not hold up against a direct comparison between groups. Of course, when the p-value of the mutant group is presented and shown to be very close to .05, the logical flaw becomes more evident and many scientists would interpret their own findings with caution, even if using EWOCs. But consider alternate scenarios where wild-type mice were perhaps /x.04 and mutants were *p<.12* (Figure 1C). Often a result of *p<.*12 would not be considered to even be approaching significance and would not be shown. Yet this result could equally fairly be stated as “We are 96% certain that the wild-type mice are social, and 88% certain that the mutant mice are social.” Expressed this way, few scientists would be confident that the mutant mice have a significant social deficit.

It could be argued that sociability in this task should be considered a binary outcome measure rather than a quantitative trait. Yet, evidence suggests this is not a categorical phenotype and that these data are indeed continuous. Multiple studies have now shown that typical sociability can be heightened or increased following stimulation of different pathways in the brain (Shin et al., 2018; Walsh et al., 2018). For example, optogenetic stimulation of the dorsal raphe neurons or their fibers in the nucleus accumbens increased the social preference index in WT mice (Walsh et al., 2018). Clearly this phenotype has a range that can be altered and deserves appropriate quantification. We have tried to make the argument here that directly comparing groups using an index, such as a social preference score, creates a suitably quantitative design, provided sufficient *n* is utilized, to overcome variability inherent in mouse behavior.

Furthermore, we have included power analyses to help guide the selection of sample sizes that will be needed to confidently overcome this variability. These sample sizes also assume a complete loss of sociability in the mutants. If the phenotype is only partial, sample size would have to be correspondingly higher. Nonetheless, while the sample size required in C57BL6/J is substantially higher than often utilized (Figure 2A), it is still reasonably achievable. However, the very high sample size required in some combinations of sex and strain suggests that considering new variations of the method that further automate the task, or that collect more repeated measures of the same mice to reduce the per mouse variance, could offer pragmatic solutions to improving power. Indeed, it is interesting that the *social novelty* trial is better powered (because of its larger effect size) than the *sociability* trial. It might be that further exposing the same mice to the Social Approach Task over multiple days might allow better estimates of the social preference of each, enabling studies that don’t require as large a sample size.

In our review of studies investigating High Confidence ASD genes, almost half of studies we examined used a flawed statistical logic to interpret the Social Approach Task results. Of these studies, 85% (11/13) concluded that the mutation impaired social behavior, and it is worrying that a substantial fraction of these might be false positives. Yet, despite the flawed statistical approach, it is possible these studies would truly show a difference between mutant and controls if the data were analyzed with an appropriate between-subjects design. For the authors with the primary data, it may be worth assessing whether this is the case. For example, in one of our prior publications, along with the standard paradigm, we employed a variation of the task we hypothesized might be more sensitive to measure preference for social novelty (cagemate versus novel conspecific) (Dougherty et al., 2013). We also examined time spent investigating a cagemate versus an empty cup. We encountered an odd situation in which the mutant mice showed a significant preference for the cagemate, whereas the control mice did not. We interpreted these within-subject differences as no deficits in sociability towards a cagemate in the mutant mice given that there were no between-subjects differences in time with the cagemate or empty cups. However, while we conducted a full repeated measures ANOVA design that included between-group simple main effects, we did not provide those results and explicitly state that the between-subjects comparisons were non-significant, thus creating ambiguity in the interpretation of our results. Therefore, here we conducted a reanalysis of these data using the preference score. This provides clear evidence that there was no difference between genotypes for sociability towards a cagemate (Control *M=*55.48, SD=9.96; Mutants *M=*62.72, SD=13.38; *t*(16)=1.226, *p*=.238). We provide this example of our own data to demonstrate how ambiguous studies can be quickly reanalyzed for clarity. Key studies that used EWOCs may benefit from corrigendums or preprint postings clarifying what the results are when the data is reanalyzed using direct statistical comparisons between groups. If prior studies were actually not significant, it could have important implications on future studies involving these ASD liability genes.

Clearly, within-subjects analyses can still have an important place in the analytical approach to Social Approach Task data as a control analysis and/or post hoc test. For example, control-level preferences are certainly useful to confirm validity and success of the experiment. Without confirmation of a preference for time spent with the social stimulus cup versus the empty/novel object cup in the control group, a direct comparison of a social preference index between controls and the experimental group may be meaningless; without a significant preference among control animals, there is no standard performance against which to compare the experimental performance. Indeed, the absolute time values of both groups are also important to examine during data analysis. There may be an instance in which the social preference index is not different between groups, but the absolute time spent with the stimuli is greatly reduced or increased in the experimental group. A great example of this can be found in Lee *et al.* (2015), in which the greatly reduced investigation times in *Shank2* homozygous mutants was found to be due to motor stereotypies. This interesting phenotype may not have been elucidated if only the social preference index was examined. Visual investigations of absolute time plots and additional analysis with a repeated measure ANOVA should always be part of the analytical pipeline of these data.

While we have highlighted the occurrence of EWOCs with regards to this one assay, this flaw certainly has been seen in a variety of other experiments in the past (Nieuwenhuis et al., 2011), and the same erroneous logic could easily be applied to a variety of other experiments in behavior (e.g. lack of alternations in T-maze) and beyond. We have been very deliberate in developing a novel term as we hope that providing a simple name for the phenomena (“EWOCs“) will aid in rapid recognition of this flaw when it occurs. More importantly, we hope the presentation of a simple solution (direct statistical comparisons) will encourage authors, editors, and reviewers to root out this kind of inference generally from the literature, and from this assay specifically.

Excellent standardized behavioral assays are essential for assessing face validity of mouse models of ASD liability and discovering new therapeutic options. A vital aspect of the validity and reliability of an assay is appropriate interpretation of the data, which requires the correct statistical approaches. The Social Approach Task is a valuable tool to assess mouse social approach behavior, one domain that could be related to the abnormal social phenotype in ASD. As such, it has been used extensively over the last 14 years and will likely continue to be frequently applied to various mouse models. Our hope, moving forward, is to begin to apply more appropriate statistical analyses to the Social Approach Task data so that accurate, reliable, and reproducible conclusions are drawn across ASD liability models. This will allow the ASD research community to move forward confidently with studies of new therapeutic strategies based on convincing and concrete results.

## Acknowledgements

This work was supported by 1R01MH107515 (J.D.D.), NIH training grant 5T32GM007067-43 (K.R.N.) and NSF Graduate Research Fellowship DGE-1745038 (K.R.N.).

